# A 3D bioprintable hydrogel with tuneable stiffness for exploring cells encapsulated in matrices of differing stiffnesses

**DOI:** 10.1101/2022.10.06.511222

**Authors:** Eric Y. Du, MoonSun Jung, Joanna Skhinas, Maria K. Tolentino, Niloufar Jamshidi, Jacinta Houng, Kristel C. Tjandra, Martin Engel, Rob Utama, Richard Tilley, Maria Kavallaris, J. Justin Gooding

## Abstract

*In vitro* cell models have undergone a shift from 2D models on glass slides to 3D models that better reflect the native 3D microenvironment. 3D bioprinting promises to progress the field by allowing the high throughput production of reproducible cell-laden structures with high fidelity. As this technology is relatively new, the current stiffness range of printable matrices surrounding the cells that mimics the extracellular matrix environment remains limited. The work presented here aims to expand the range of stiffnesses by utilising a 4-armed polyethylene glycol with maleimide functionalised arms. The complementary crosslinkers comprised a matrix metalloprotease (MMP)-degradable peptide and a 4-armed thiolated polymer which were adjusted in ratio to tune the stiffness. The modularity of this system allows for a simple method of controlling stiffness and the addition of biological motifs. The application of this system in drop-on-demand printing is validated in this work using MCF-7 cells which were monitored for viability and proliferation. This study shows the potential of this system for the high-throughput investigation of the effects of stiffness and biological motif compositions in relation to cell behaviours.

## Introduction

Questions pertaining to cell biology have been traditionally explored using two-dimensional (2D) technologies with cells cultured on top of a biocompatible surface. While these 2D cultures are the very bedrock of the entire field of cell biology, the literature is showing a progressive shift towards the more common use of 3D cultures whereby cells are encapsulated inside a matrix.^1, 2^ Cell properties such as proliferation and migration have been reported to differ in 2D and 3D environments.^3, 4^ The shift to 3D cell cultures has partly been motivated by commercially available hydrogels such as Matrigel, a basement-membrane matrix extracted from Engelbreth–Holm–Swarm mouse sarcomas,^5^ as mimics of the extracellular environment (ECM). The importance of the results obtained with Matrigel have motivated cell biologists to search for ECM mimics with lower batch variability, greater biochemical complexity, and better control over their physical and biological properties.^5, 6^ With the advances and progress made in ECM mimics, it is becoming clear that expanding the range of ECM mimics available will greatly benefit the field as would high-throughput ways of integrating these materials into 3D cell cultures.

Bioprinting is a technology that has opened up the avenue for creating complex 3D cell-laden constructs with high fidelity and reproducibility. This technology has therefore been applied to many biomedical fields such as tissue engineering, drug delivery, and cancer research.^7–9^ A wide array of printing techniques for delivering cells with ECM mimics and scaffolds have become available to researchers to create structures for biomedical applications.^10–12^ Most of these methods use extrusion type printing. However, for producing 3D cell cultures, drop-on-demand bioprinting, where a droplet of liquid bioink is followed by a droplet of activator solution to trigger gelation, shows great promise. This is because the drop-on-demand printing method offers high throughput, high resolution, uniform, and reproducible printing of structures and minimal cell damage.^13^ Therefore, in principle, different cell types and bioactive ingredients can be precisely positioned in a multicellular structure.^13^ Drop-on-demand printing however imposes strict criteria that the ECM material must satisfy. Most notably, the gel must be formed from two separate liquids with relatively low viscosity. Secondly, they must gel almost instantaneously such that by the time a droplet is dispenses the previous droplet has begun to solidify into a hydrogel. Although some synthetic and biopolymer ECM mimics achieve these criteria their stiffnesses are low and only allow recapitulation of soft tissues.^14–16^ Even commonly used synthetic matrices such as Matrigel or collagen can only produce low stiffness values. Therefore, as stiffer printable hydrogels are required to mimic a larger range of tissues.

In general, increasing the concentration of a polymer will increase the stiffness of the resulting hydrogel to some extent.^17, 18^ However, higher concentrations of polymers are correlated with higher viscosity.^19, 20^ As such, this approach inevitably leads to a detrimental effect on the ability to print using drop-on-demand printing as the solution becomes more viscous and is no longer able to be ejected from the nozzles. We previously demonstrated that a polyethylene glycol polymer with 4 maleimide functionalised arms (PEG-4-Mal) and a bis-thiol crosslinker can be bioprinted using a drop-on-demand bioprinter and utilised to form 3D cell models.^21^ That bioink could achieve a stiffness of 2.7 kPa, which equates to a stiffness similar to malignant breast tissue, at 20% w/v polymer which was the maximum achievable. Therefore, in designing a printable matrix for a drop-on-demand printer, an alternative approach to increasing the concentration of polymers to increase the stiffness is required.

The purpose of this paper is to develop a printable cell laden matrix using the PEG-4-Mal hydrogel system that is easily translatable to commercial applications. Gold standards in commercially available ECM mimics such as Matrigel or collagen are relatively soft, reaching stiffness values of 0.05-1 kPa.^22–25^ This work describes a 3D printable network with a tuneable stiffness range of 1.9-6 kPa. The base ECM mimic system used herein is based on the bioink utilises a PEG-4-Mal bioink and a 2-armed matrix metalloproteinase (MMP) cleavable crosslinker. Different crosslinkers change the stiffness of a network due to differing degrees of conformational freedom in the crosslinking species. This is the strategy adopted in this work as the stiffness of a network can be adjusted with relatively little change to the viscosity of the polymer solution. By using a 10 kDa 4-armed PEG-4-Thiol crosslinker to replace some of the 2-armed crosslinker, a stiffer network is achieved with little change to the printability of the bioinks. The stiffness of the hydrogels was adjusted by changing the ratios of polymer crosslinker to peptide crosslinker. We demonstrated that the system allows for the generation of printable PEG-4-Mal hydrogels ranging from 1.9 kPa to 6 kPa, which encompass tissues such as liver and pulmonary valve tissues.

## Experimental

### Bioinks and Activators

PEG-4-Mal and MMP cleavable crosslinkers to make base bioinks and their respective activators were purchased from JenKem and Genscript. The 10 kDa 4-armed polyethylene glycol with thiol functionalisation (PEG-4-Thiol) was purchased from JenKem. The Peg-4-Mal was prepared at either 16% w/v or 20% w/v in phosphate buffered saline (PBS). The MMP-cleavable crosslinker was then prepared at a 2:1 molar ratio to the PEG-4-Mal and dissolved in PBS. To prepare the stiffer gels, 200 mg of PEG-4-Thiol was dissolved in 1 mL of PBS. All solutions were then sterile filtered using a 0.22 ⍰m filter The MMP cleavable crosslinker and PEG-4-Thiol solutions were combined in 3:1 and 1:1 MMP cleavable crosslinker:PEG-4-Thiol volumetric ratios to create the activators used for the stiffened gels. The bioink is unchanged for the stiffened gel compositions.

The Rastrum bioprinter was controlled via a combination of the proprietary Rastrum software. Parameters such as opening time, pressure, and droplet placement is all adjusted in the software. PEG-4-Mal was printed at a pressure range of 32-70 kPa and with nozzle opening time ranges of 320-480 ms. The activators containing PEG-4-Thiol and MMP-cl were printed using a pressure range of 32-80 kPa and an opening time range of 320-480 ms.

### Rheology

Rheology measurements were performed in triplicate for each condition. Samples were printed as cylinders 1 mm high and 25 mm in diameter. Samples were swelled overnight in phosphate buffered saline and measured using an Anton Paar MCR302 Rheometer using a 25 mm measuring plate setup. Frequency was kept at a constant rate of 1 Hz and the strain was kept at a constant 0.1%. Temperature was set to 37 °C. Storage modulus was taken as the stiffness of the hydrogel.

### Cryo-scanning electron microscope

Samples for cryo-scanning electron microscopy were printed as a cylinder 10 mm in diameter and 5 mm in height. Each condition was performed in triplicate. In a typical cryo-SEM experiment, a piece of hydrogel was transferred to a specimen holder and submerged in liquid nitrogen for 30 seconds. The sample was fractured with a cooled blade and transferred to a specimen stage in the vacuum stage of a Cryo-SEM microscope. The morphology of the pores was then assessed using a JEOL JSM-6490LV SEM at the Electron Microscopy Centre at the University of Wollongong. The SEM was operated at 15 kV accelerating voltage and a spot size of 16 (~200 pA) with a WD of 15 mm. The imaging was performed in a secondary electron imaging mode using Oxford Instruments AZtec. Imaging was done with frame integration from 8 – 24 frames and dwell times of 1 – 5 μs.

### Pore analysis

Pore size and shape were measured using an ImageJ algorithm. In brief, the foreground and background were separated by local intensity thresholding to create a binary mask. The voids of the pores detected by the binary masks were then measured via the analyse particle function. Sizes were adjusted based on the scale bars. Objects that were detected via the binary mask but which were too small (consisting of single pixels) or too large (clumped pores with boundaries indistinguishable via thresholding) were filtered out. Parameters for filtering were: ; pore area <3 ⍰m; pore area <10 ⍰m2 and roundness >0.9.

### Cell culture

MCF7 were purchased from ATCC and cultured in DMEM media supplemented with 10 % FCS. Cells were maintained in a humidified atmosphere containing 5 % CO2 at 37 °C and were mycoplasma free. The cell line was authenticated using short tandem repeat profiling at Kinghorn Centre for Clinical Genomics, Australia.

### Live and dead cell staining

Cell viability analysis was performed using Live/Dead viability/cytotoxicity kit, for mammalian cells (Invitrogen, Cat No. L3224). Briefly, cells were bioprinted in multi-well plates and cultured for 7 days post-printing. At day 7, cells were rinsed with DPBS and stained with 100 μL of live/dead stock solutions (10 μM ethidium homodimer-1 (EthD1) and 5 μM calcein AM in DPBS) and incubated for 30 min. Images were taken at 5X magnification using green fluorescence channel for live cells and red fluorescence channel for dead cells using CellDiscoverer 7 (Zeiss). Images were analysed and visualized using Arivis 4D software.

### Statistical analysis

Statistical analyses were performed with GraphPad Prism 9.0. For comparison of multiple samples, Kruskal–Wallis one-way analysis was used.

## Results and discussion

### 3D printable hydrogel with tuneable stiffness

A hydrogel system is used in this work where a maleimide-thiol reaction between a 10 kDa polyethylene glycol with PEG-4-Mal and a matrix-metalloproteinase (MMP) cleavable peptide sequence flanked by thiol groups. The cell-adhesive peptide, RGD (Arg-Gly-Asp), was conjugated to the PEG-4-Mal to explore the effects of introducing a biological motif to the system (Fig. 1). This is achieved by the addition of 2 mM CRGDS to the PEG-4-Mal prior to crosslinking. A polymer crosslinker in the form of a 10 kDa 4-arm polyethylene glycol with thiols on each arm was used in combination with the peptide crosslinker to form stiffer hydrogels. The network formed by using the polymer crosslinker was shown to be stiffer than the peptide crosslinker as it potentially forms four crosslinks, rather than only a possible two with the peptide linker, which limits the conformational freedom of the final polymer network (Fig 1A). Structures were printed using a drop-on-demand printer which uses two axes to control x and y positioning while building structures in the z direction is achieved by printing layer upon layer (Fig. 1B).

**Figure 1.**
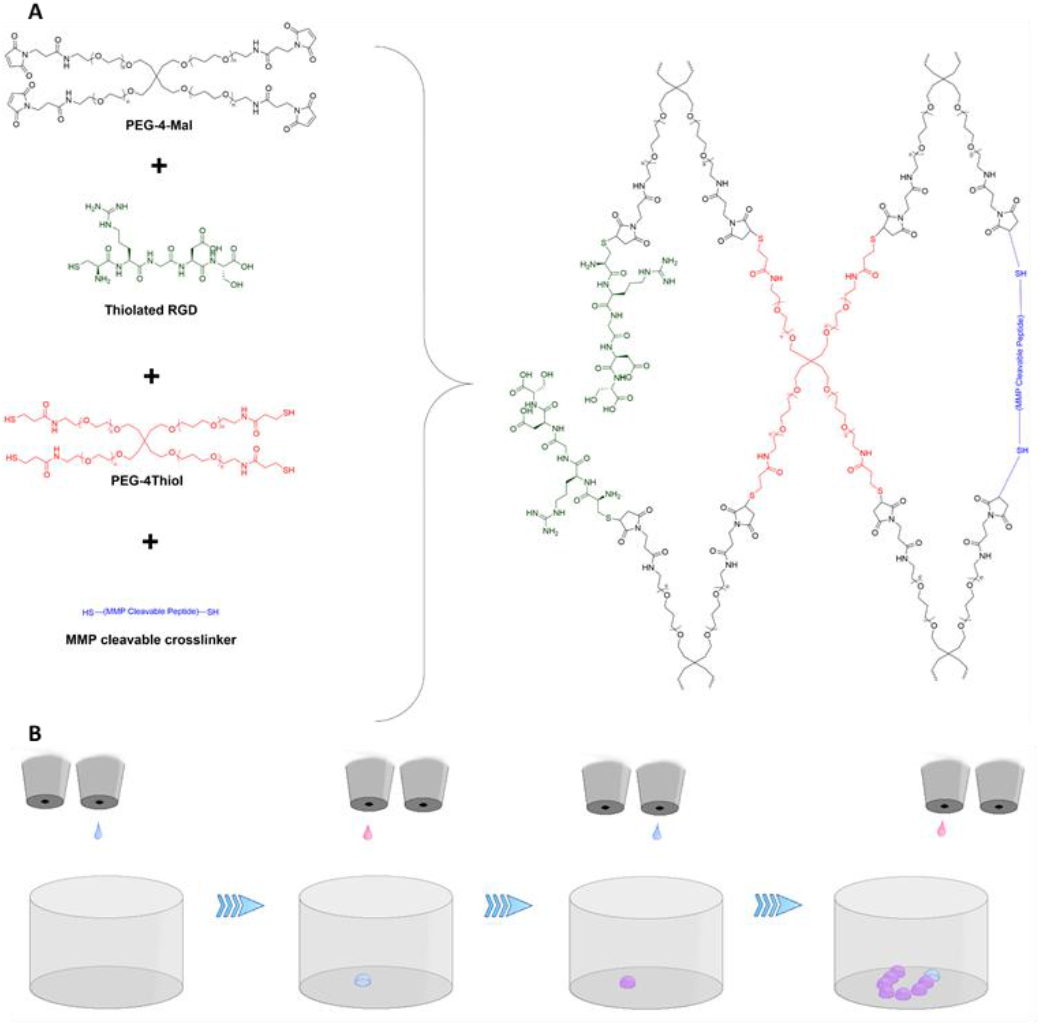
The schematic of the polymer and printing systems. (A) The chemical crosslinking mechanism with the MMP cleavable crosslinker, the stiffening crosslinker, and the thiolated RGD. (B) The drop-on-demand printing strategy.

Storage moduli (G’) values from oscillatory rheology were used as the indicator of hydrogel stiffness. The base system made with 10 kDa PEG-4-Mal with the peptide cross-linker, but without the 4-arm polymer crosslinker to increase the stiffness, gave a storage modulus (G’) value of 1.9 kPa at a concentration of 16% w/v with no PEG-4-Thiol crosslinker (16PEG0) and a G’ of 2.5 kPa at 20% w/v with no PEG-4-Thiol crosslinker (20PEG0) as shown in Figure 2 and Table 1. Increasing the amount of 4-arm polymer cross-linker to a molar ratios of 6:1 and then 2:1 peptide:polymer crosslinker resulted in stiffer hydrogel ECM mimics. The hydrogel made with 6:1 peptide:PEG-4-Thiol at 20% w/v (20PEG14) produced a higher G’ of 4 kPa. By increasing the ratio to 2:1 peptide:PEG-4-Thiol (20PEG33), the printed hydrogel exhibited a G’ of 6 kPa (Fig. 2 and Table 1). The 4 polymer systems together have a stiffness gradient from 1.9 kPa to 6 kPa which spans the ranges from lung, liver and heart tissue.^26–28^ A second set of gels was also investigated that incorporates an RGD motif by reacting a CRGDS peptide sequence to ~8% of the maleimide arms of the PEG-4-Mal prior to mixing the polymer with the crosslinkers. The addition of RGD in conjunction with the ability to tune the stiffness, adds an extra degree of biomimicry as RGD is a prevalent sequence found in the extracellular matrix and this can be used to explore the combined effects of RGD and stiffness on cells. The same method of stiffening was adopted to modify the stiffness of the RGD bearing peptide using 10 kDa PEG-4-Mal with RGD at 16% w/v (16PEG0-RGD) and at 20% w/v (20PEG0-RGD). PEG0-RGD was added to either 6:1 or 2:1 peptide:PEG-4-Thiol crosslinker respectively to give PEG14-RGD and PEG33-RGD which gave a stiffness gradient from 1.2 to 5 kPa. Noticeably, the addition of RGD decreased the stiffness slightly, compared to when no RGD was used, as the conjugation of the RGD utilises some of the maleimide groups and therefore reduces the overall extent of crosslinking.

**Table 1.** A summary of the storage moduli (G’) data of modified hydrogels.

| Condition | $G'$ (kPa) |
| --- | --- |
| 16PEG0 | 1.9 |
| 20PEG0 | 2.7 |
| 20PEG14 | 3.9 |
| 20PEG33 | 6.0 |
| 16PEG0-RGD | 1.5 |
| 20PEG0-RGD | 2.2 |
| 20PEG14-RGD | 3.7 |
| 20PEG33-RGD | 5.0 |

**Figure 2.**
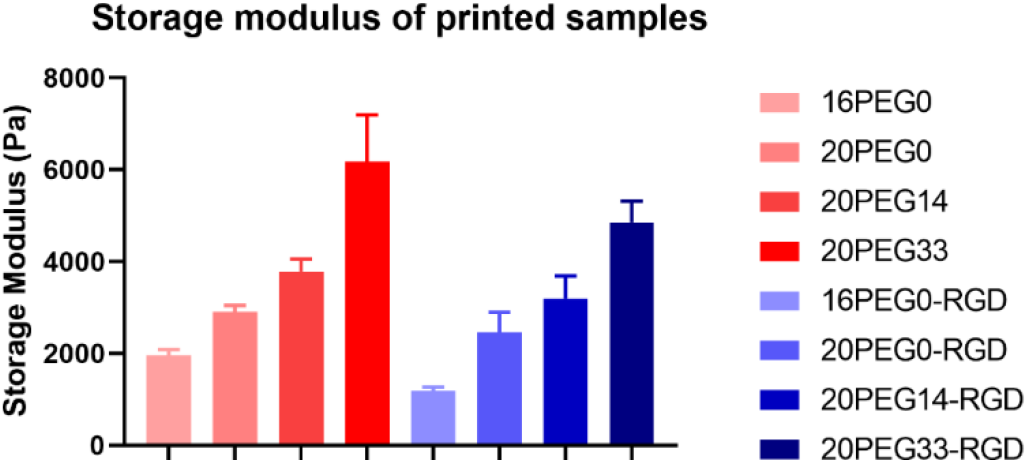
The stiffness characterisation represented as storage moduli (G’) of the printed hydrogel samples assessed by rheology.

As the stiffness of a hydrogel is related to the underlying hydrogel structure, the pore size and shape of the hydrogels were analysed using cryo-SEM to correlate any changes in stiffness to the pore network (Fig. 3). Increasing the concentration of PEG from 16% w/v in 16PEG0 to 20% w/v in 20PEG0 resulted in a decrease in pore size. Pore area decreased from 28 μm2 to 13 μm2 while perimeter decreased from 24 μm to 17 μm (Table 2). Further modification of the system by the replacement of some of the peptide with the thiol-based polymeric cross-linker, without the addition of RGD, did not change either the pore area or the pore perimeter significantly. Pore shape was also investigated using shape roundness and solidity. Roundness is a numerical representation of the gross shape and compares the minor axis to the major axis. Solidity examines the edges of a shape and compares the area of the shape to the area of a convex hull. The convex hull is defined as the smallest convex shape that can be drawn by connecting the outermost corners of a shape. While the solidity remains unchanged between the three gels, the roundness of the 16PEG0 condition is higher than other conditions. This indicates that in the softest condition, the overall minor axes are becoming more similar to the major axes and becoming rounder. This matches the pore size data as this suggests that the smaller pores of the stiffer conditions are more compactly pressed together causing the pores to be less round. The same pore analysis was performed on the gels containing RGD to assess the changes in the underlying network as a result of peptide conjugation. It was found that in hydrogels with RGD, the stiffer samples had larger pore areas while the size of the pore perimeter showed no obvious trends (Fig. 3b). Solidity and roundness were also measured which shows that the solidity of the softest condition, 16PEG0-RGD, is lower than all other conditions which suggests that the edges of the pores are more tessellated in the stiffer samples.

**Table 2.** Summary of pore size and shape as analysed from Cryo-SEM images of printed hydrogel samples. Performed for n=3

| Condition | Area (μm<sup>2</sup>) | Perimeter (μm) |
| --- | --- | --- |
| 16PEG0 | 28.0 | 24.2 |
| 20PEG0 | 13.5 | 17.1 |
| 20PEG14 | 13.3 | 17.3 |
| 20PEG33 | 14.1 | 18.3 |
| 16PEG0-RGD | 13.6 | 26.2 |
| 20PEG0-RGD | 16.3 | 19.6 |
| 20PEG14-RGD | 21.8 | 20.4 |
| 20PEG33-RGD | 24.5 | 22.3 |

**Figure 3.**
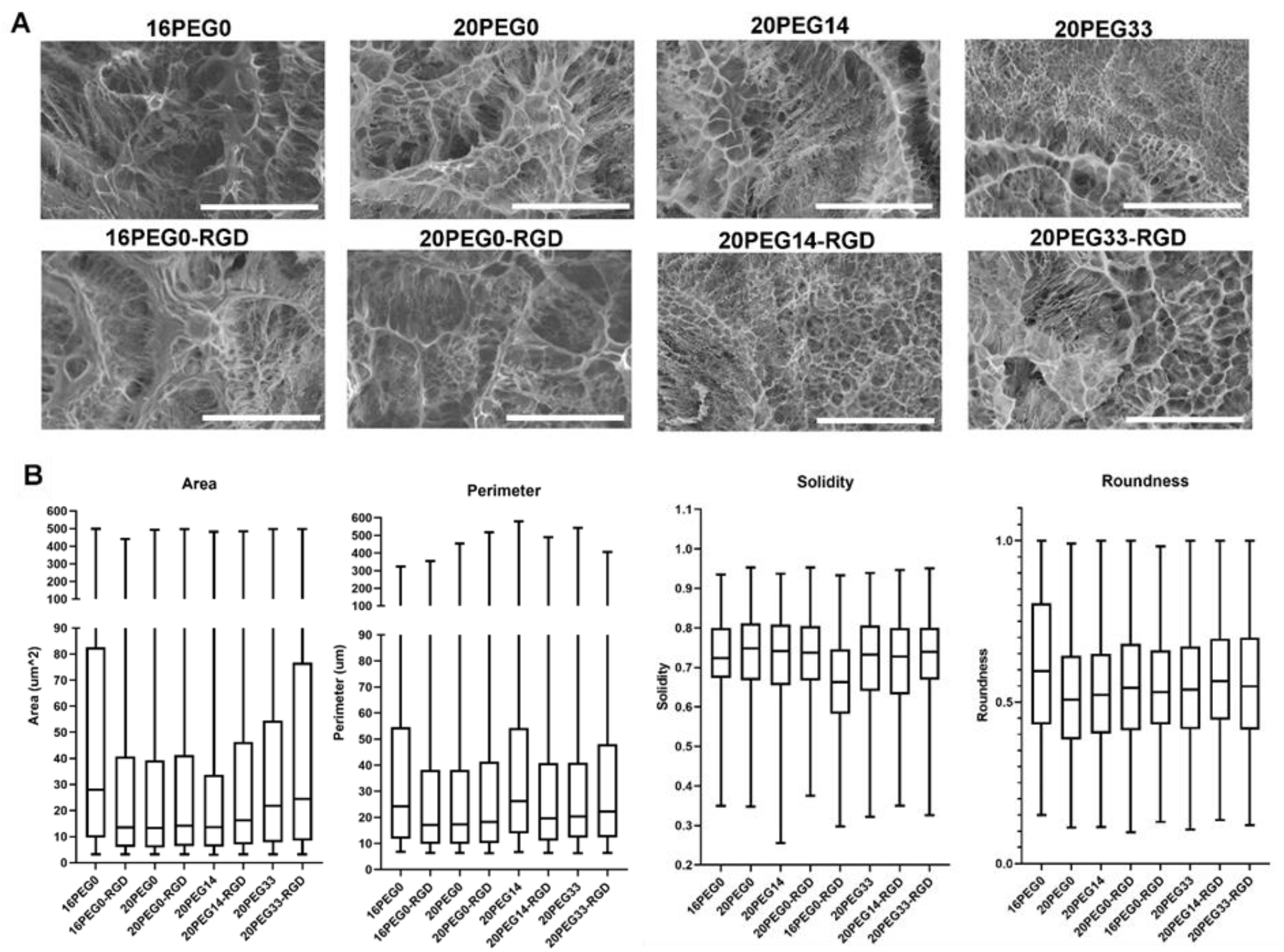
Pore characterisation assessed by Cryo-SEM. (A) The representative Cryo-SEM images of 20% w/v 10 kDa with and without RGD with increasing amounts of stiffness with all scale bars at 100 μm. (B) The analysis of the pore size and shape of printed bioink samples. Size is assessed using area and perimeter. Shape is assessed using solidity and roundness. Performed for n=3.

### Cell viability and proliferation

To determine the cytocompatibility of the bioinks for use in bioprinting structures, MCF-7 breast cancer cell line was printed with each hydrogel condition and observed to grow steadily over the 7-day period in all conditions. Cell metabolic activity was assessed via an AlamarBlue Assay (Fig. 4A). The stiffer conditions were observed to have lower amounts of growth which indicates that the cells prefer to grow in softer gel conditions. This is likely due to MCF-7 cells being derived from breast tissue which is a soft tissue. Malignant breast tumours have been reported to have a Young’s modulus of 2.5 kPa. ^29, 30^ This corresponds to a storage modulus value of approximately 1 kPa, as calculated by a previously reported formula, and is closest in stiffness to the softest bioinks printed (16PEG0).^26^ Therefore, these cells are accustomed to growing in a natively soft environment rather than the stiffer hydrogels, resulting in slower growth. In addition, LIVE/DEAD staining and phase contrast imaging confirmed that MCF-7 cells remained highly viable in all hydrogel conditions at 7 days post-printing (Fig. 4B). This indicates that the addition of the stiffening agent does not negatively impact the viability of the cells and that the cells can survive the denser crosslinking mechanism.

**Figure 4.**
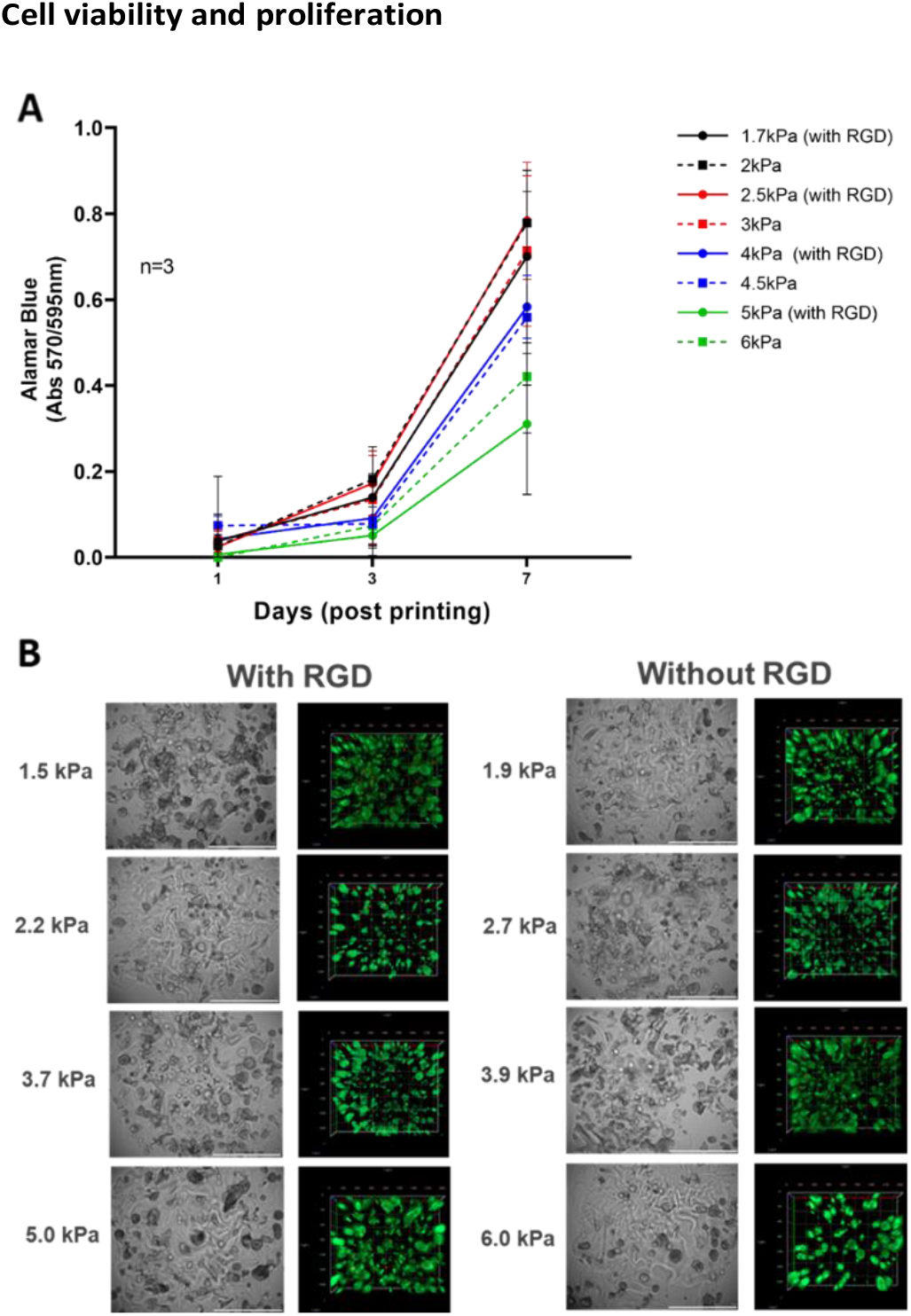
Cell viability as determined by (A) an AlamarBlue assay and (B) a LIVE/DEAD stain which show that cells are alive and growing over a period of 7 days. Phase contrast images were used as representative images were also taken, and all cells successfully form spheroids after 7 days indicating normal cell behaviour. All scale bars are 100 ⍰m. Performed across n=3.

The capability of the drop-on-demand bioinks were also demonstrated by using the bioinks to explore the formation of spheroids in printed matrices of differing stiffnesses. MCF-7 cells were printed with the bioinks and incubated over 7 days, at which point the spheroids were imaged and analysed for size. As seen in Figure 5, spheroid sizes remained similar across all ranges except in the 20PEG33-RGD condition which showed significantly smaller spheroids. This is in line with the proliferation data which also shows that the 20PEG33-RGD condition had the lowest proliferation of MCF-7 cells. The number of spheroids also decreased with increasing stiffness which follows the trend in the proliferation data in Figure 4 which emphasises the stiffness being an important factor in the regulation of MCF-7 cells. These findings are in agreement with work in literature which has shown that stiffer matrices hinder spheroid formation due to greater compaction of the 3D matrix.^31–33^ The data out-put demonstrated here highlights the capability of the bioink in conjunction with a compatible bioprinting platform for the high-throughput study of cells.

**Figure 5.**
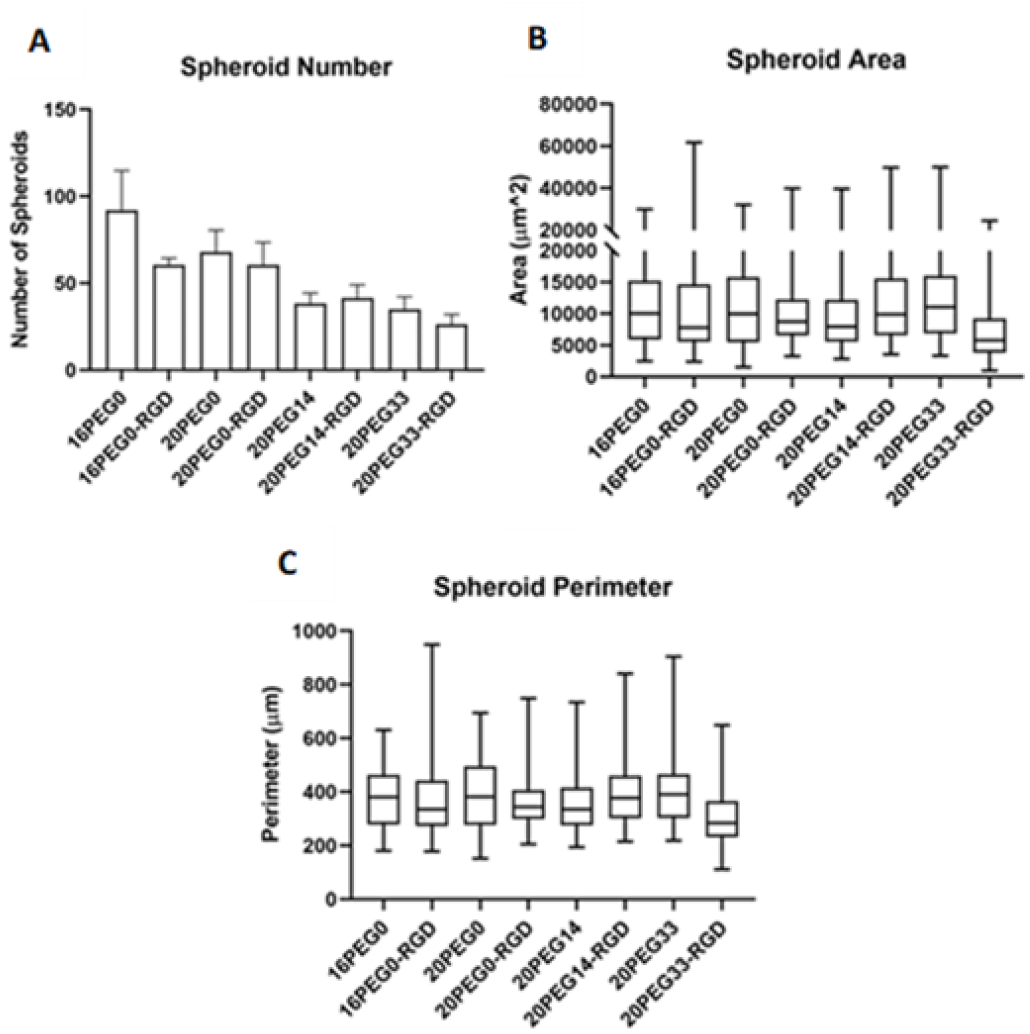
Data for spheroid (A) number, (B) area, and (C) perimeter with MCF-7 cells in printed hydrogels. Cells were cultured over 7 days. Data analysed with ImageJ from images taken with confocal microscopy. Performed for n=3.

## Conclusions

In this work, we have printed a stiff cell-laden matrix using a hydrogel formulation that has never been printed before in a drop-on-demand printer. It is shown that these cell matrices are considerably stiffer than those previously printed with this technology expanding the range more than 2-fold from 1.9 kPa to 6 kPa. Furthermore, these printed ECM mimics are significantly stiffer than the currently available commercial gold standards such as Matrigel or collagen which reach stiffnesses of 1 kPa.^22–25^ We also show that the stiffness and composition can be easily tuned by adjusting the amount of PEG-4-Thiol or RGD in the crosslinker. The possibility of studying cells using printed structures was demonstrated by using MCF-7 in hydrogels of varying stiffness to study proliferation and viability. While cells remained viable in all conditions, in stiffer networks, the MCF-7 cells exhibited slower growth. In printing cell laden matrices, it has been demonstrated that the hydrogel matrices presented in this work are biocompatible and can be used to study cells with high fidelity. The work presented describes an simple method to create stiff cell laden hydrogels which is easily translatable to many cell types and has shown that these hydrogels can be used to control cell kinetics which will undoubtedly assist in developing and expanding high throughput biological assays.

## Author Contributions

Eric Y. Du (conceptualisation, methodology, validation, investigation, formal analysis, writing – original draft, writing – review and editing, visualisation), MoonSun Jung (methodology, validation, investigation, formal analysis, writing – review and editing), Joanna Skhinas (methodology, validation, investigation, formal analysis, writing – review and editing), Maria K. Tolentino (software, data curation, formal analysis), Niloufar Jamshidi (software, data curation, formal analysis), Jacinta Houng (formal analysis), Kristel C. Tjandra (methodology, formal analysis), Martin Engel (validation, writing – review and editing), Rob Utama (validation, writing – review and editing), Richard Tilley (supervision), Maria Kavallaris (supervision, project administration, funding acquisition), J. Justin Gooding (conceptualisation, supervision, project administration, funding acquisition, writing – original draft, writing – review and editing, visualisation)

## Conflicts of interest

The authors declare that there are no known conflicts of interest, whether financial or personal, that have affected the work reported in this article.

## Acknowledgements

The research was financed by the Australian Research Council Linkage Program (LP170100623), the National Health and Medical Research Council Investigator Grant (GNT1196648), the National Health and Medical Research Council Principal Fellowship (APP1119152), the University of New South Wales School of Chemistry, and the Children’s Cancer Institute (affiliated to the University of New South Wales). The authors also thank the Katharina Gaus Light Microscopy Facility and the University of Wollongong Electron Microscopy Centre for their support and resources involved in this work.

## Notes

### Competing Interest Statement

The authors have declared no competing interest.

